# Model-based intensification of CHO cell cultures: one-step strategy from fed-batch to perfusion

**DOI:** 10.1101/2022.05.19.492635

**Authors:** Anne Richelle, Brandon Corbett, Piyush Agarwal, Anton Vernersson, Johan Trygg, Chris McCready

## Abstract

There is a growing interest in continuous processing of the biopharmaceutical industry. However, the technology transfer from traditional batch-based processes is considered a challenge as protocol and tools still remain to be established for their usage at the manufacturing scale. Here, we present a model-based approach to design optimized perfusion cultures of CHO cells using only the knowledge captured during small-scale fed-batch experiments. The novelty of the proposed model lies in the simplicity of its structure. Thanks to the introduction of a new catch-all variable representing a bulk of by-products secreted by the cells during their cultivation, the model was able to successfully predict cellular behavior under different operating modes without changes in its formalism. To our knowledge, this is the first experimentally validated model capable, with a single set of parameters, to capture culture dynamic under different operating modes and at different scales.

## 1 Introduction

Currently, only one out of every 10,000 new drug candidates reaches the market. It takes on average 10 years from the discovery of a drug compound until its approval by federal agencies. The probability of clinical success is less than 10% (from Phase 1 to launch)^1,2^. As a consequence, the cost of drug development is constantly increasing, with a current annual expenditure of more than 2 billion euros, while the actual revenues do not follow the same trend^3,4^.

In this context, we observe that biopharmaceutical companies tend to outsource their early activities in order to reduce their costs and to be more agile and more flexible around potential market disruption^5,6^. Their main focus becoming then the go-to market activities (process development and product manufacturing). This means that they have growing need to accelerate their operational tasks. The path to acceleration for biopharmaceutical industry relies mostly on digitalization of process information to fasten process development and intensification of operations to increase productivity and enable more flexibility in the production.

Digital bioprocessing is expected to provide significant competitive advantage to industry adopters (e.g., rapid process prototyping, improved process performance and product quality, and de-risked transfer to manufacturing)^7^. This digital transformation relies on the computerization of the information used and generated at each step of a product development process. Once all the information is digitized, it needs to be accessible, organized and contextualized (i.e., data management). This structured digital information can therefore be used to feed data analytics and associated modeling tools to generate valuable insights for further process optimization and control, thereby fasten process development^8,9^.

Today, the industry standard for proteins production such as monoclonal antibodies is a fed-batch process. However, the productivity of such a process can be significantly improved by implementing a continuous culture strategy to intensify the volumetric productivity. Such approach can lead to an increase up to 10-fold of space-time yields, therefore leading to a reduction of production time by 30%^10^. These improvements enable opportunity for much smaller facilities with similar or larger productivity outputs, limiting the capital investment (for facilities and raw material costs) and providing manufacturing flexibility and sustainability^10,11^.

Such transition from traditional (fed)-batch to continuous manufacturing is facilitated by the emergence of various technological enablers^12^ and is encouraged by health authorities (i.e., US Food and Drug Administration). However, the adoption is relatively slow as many challenges remain. Indeed, scale-down models, decisional tools, equipment and procedures currently in place in most companies have been developed for fed-batch processes and cannot be transposed without significant changes. Therefore, this transition might be seeming a high cost and time demand investment to modify existing process development protocols^10,11,13^. In this context, advanced computational tools could be used to elucidate changes in process dynamics and assess the influence of varying operating scenarios. These in-silico tools provide testing platforms for early determination of process bottlenecks at minimum experimental costs and enable the design of advanced optimization strategies that will lead to optimal and stable operation^12,13^.

While this burden for digital transition and technology transfer has been observed in the past (and successfully overcome) in other industry sector (e.g., petrochemical companies, aeronautics), biopharma faces the additional challenge that its operation relies on complex biological systems that cannot be easily described using known first principles rules. Numerous modelling studies successfully characterizing the influence of measurable process conditions on culture dynamic exist in literature. Unfortunately, they often rely on numerous measurements not often available at manufacturing scale and/or complex modeling and optimization procedures requiring important computational expertise. Therefore, these model-based intensification strategies are difficult to be transferred at industrial scale in spite of their most likely success^13^. Here, we focused on the development of a modeling structure enabling the description of upstream bioprocess dynamics and the transfer between operations (specifically, from fed-batch to continuous culture) at different scales (from Ambr^®^ 250 to 2L) with a single set of kinetic parameters. We have demonstrated that the growth model identified using fed-batch cell cultures can be used to design intensified culture conditions in a one-step strategy. To our knowledge, this is the first experimentally validated methodology providing simulation capabilities appropriate for optimization and system configuration decisions within biopharmaceutical process development and advanced control activities.

## 2 Materials and methods

### 2.1 Cell line, inoculum development, medium and analytical methods

CHO DG44 cell line (Sartorius) expressing a monoclonal antibody (mAb, IgG1) was used. All experiments were carried out using the same chemically defined media (Sartorius) and Stock Culture Medium (SCM) for the seed train. The seed train cultures were performed in 5 steps. For the fed-batch processes, these pre culture steps were performed in (unbaffled) shake flasks. For the perfusion culture, the last pre-culture step was performed in a 2L Univessel^®^. The first and second pre culture steps were performed in SCM with 15nM MTX while the others were without MTX. Cells were seeded at 0.2×10^6^ cells/mL and split every 3 to 4 days. The incubators settings for the shake flasks were: 7.5% CO_2_, temperature at 36.8 °C, 80% humidity, 120 rpm for agitation with an orbital diameter of 50mm. Fed-batch, intensified and perfusion cultures were performed with a production medium (PM - Sartorius) and with two feed media (FMA and FMB -Sartorius). Cell growth (VCC and viability) were measured using a Cedex HiRes Cell Counter (Roche).

### 2.2 Fed-batch and intensified cultures in Ambr^®^ 250

The fed-batch and intensified cultures were performed in Ambr^®^ 250 bioreactor with a working volume of 200 mL and 210 mL respectively for fed-batch and intensified cultures. The cultures were inoculated at 0.3×10^6^ cells/mL. The feeding profiles were calculated off-line following the standard applications implemented in Sartorius. For the intensified cultures, the flow rate was adjusted daily from day 3. The culture conditions were controlled at 36.8 °C for the temperature, 855 rpm for the agitation (adjusted during culture according O_2_ demand), pH 7.1 with CO_2_, 60% of DO with O_2_ and air inlets. 30 μl of antifoam (Sigma antifoam C 2%) was automatically added every 12 hour and manually added if needed. A daily glucose bolus was performed starting on day 5 of culture if the measured glucose concentration was less than 5 g/L (stock glucose solution of 400 g/L).

### 2.3 Perfusion cultures in 2L bioreactor

Perfusion culture is a type of continuous operation where the cell concentration and the volume within the bioreactor are kept constant. Specifically, the cell culture is continuously fed with fresh medium while a cell free harvest is continuously removed to keep the culture volume constant. To this end, the harvest stream is firstly directed through a cell retention device that will separate the used media from the cells (lived and dead). The cells are then re-injected in the bioreactor while the cell-free stream is collected for further purification of the drug product. A “bleed” outflow (stream presenting the same composition as the bioreactor) is further used to maintain the culture at steady-state (i.e., maintain the concentration of cells within the bioreactor constant)^11,14^ (Figure 1).

**Figure 1.**
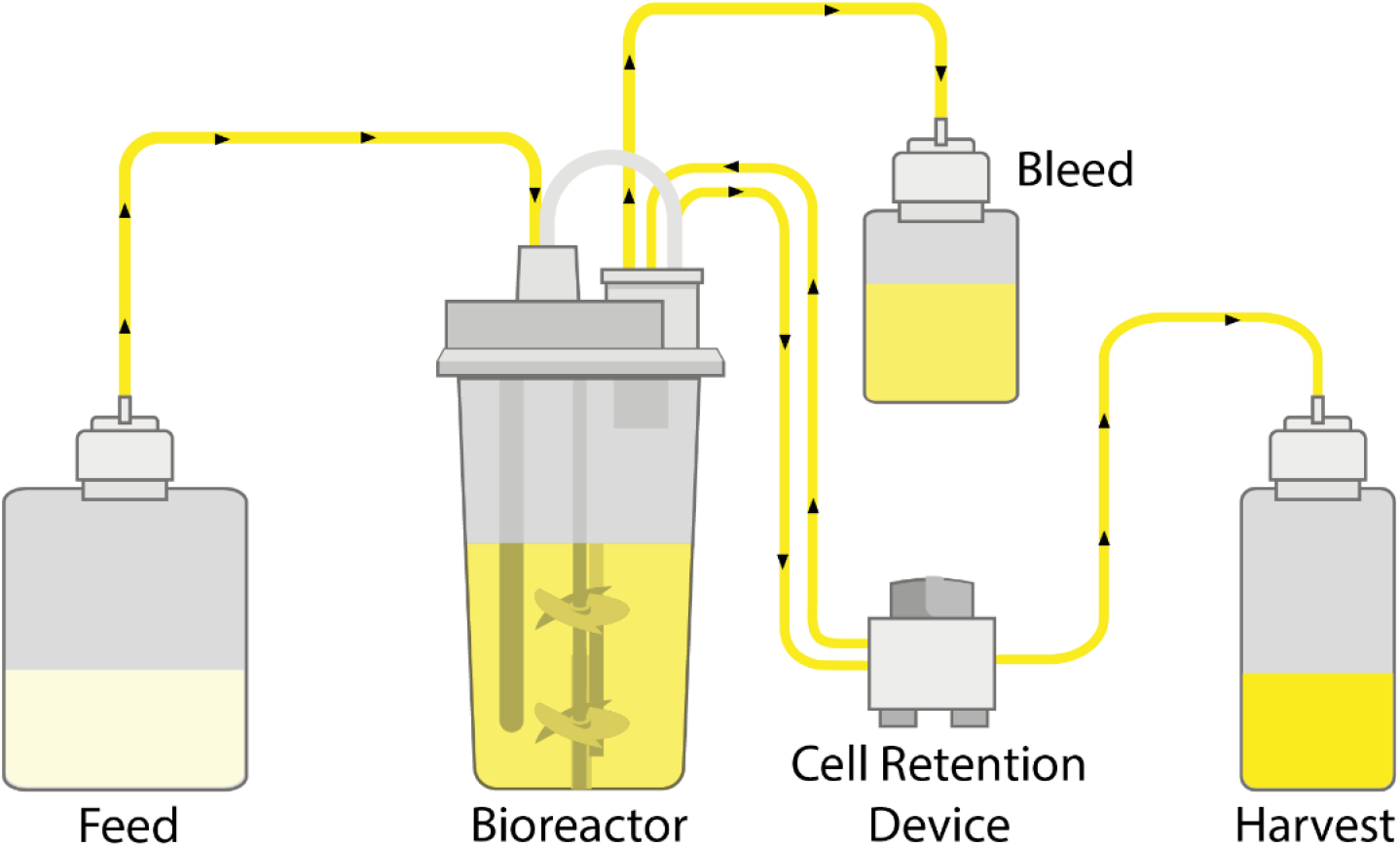
Schematic of a perfusion bioreactor. Media is continuously fed into the bioreactor (*F_F_*) and a cell free harvest is continuously removed (*F_h_*). The cell retention filter is assumed to be ideal, where only the lysed cells and other cellular by-products pass through while viable and dead cells are fed back into the bioreactor. The bleed stream (*F_b_*), containing same content as the bioreactor, is used to keep a steady concentration the cells within the bioreactor by removing cells in excess.

The rate at which the media is exchanged in this operation can be defined either by the cell specific perfusion rate (*CSPR* – media supply needed by cells by day) or by the perfusion rate (*P* - amount of bioreactor volume renewed by day). Specifically, the cell specific perfusion rate (*CSPR*) is defined as the ratio between the perfusion rate (*P*) and the viable cell density (*X_V_*) while the perfusion rate (*P*) is defined as ratio between the feeding rate *F_f_* and the volume of the bioreactor *V*):

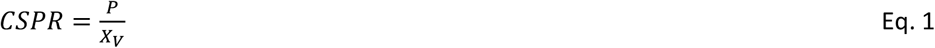

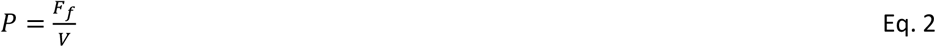

Note that contrarily of the perfusion rate, the CSPR is cell and media specific and therefore represents an important performance criterion^11,14^.

Typically, a perfusion culture is set in two phases: an intensification phase that enable the culture to growth exponentially until the target cell concentration is reached and maintained during the steady-state phase. During the intensification phase, the feeding rate (*F_f_*) is equal to the harvest rate (*F_h_*) while the bleed stream (*F_b_*) is set to zero. Using Eqs. 1 and 2, the optimal feeding rate for a given perfusion rate can be defined as follows:

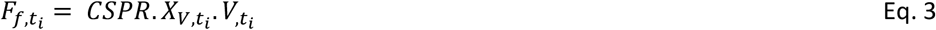

where *F_f,t_i__*, *X_V,t_i__*, *V_t_i__* are respectively the feeding rate, the viable cell concentration and the bioreactor volume at time *i*.

Once the target cell concentration (*X_V,target_*) is reached, the definition of the feeding rate (Eq. 3) can be simplified into:

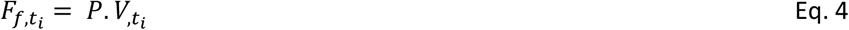

This targeted cell concentration can be maintained at steady-state by introducing the bleed stream as control variable of the process. In the context of this study, we used a Proportional-Integral (PI) controller to define the bleed rate:

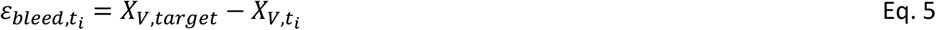

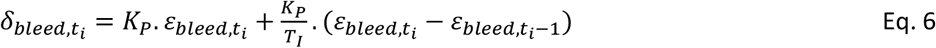

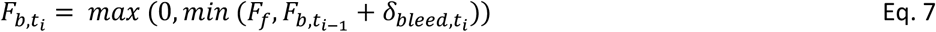

where *ε_bleed,t_i__* is the deviation at time *i* of the cell concentration (*X_V,t_i__*) from the target setpoint (*X_V,target_*), *δ_bleed,t_i__* is the controller output (with *K_P_* and *T_I_* as proportional and integral terms) that will be used to adjust the bleeding rate imposed at time *i* – 1 (*F*_*b,t*_*i*–1__) such as to maintain the target setpoint (*X_V,target_*). For this study, the PI control parameters have been hand tuned and set to *K_P_* = –0.2 and *T_I_* = 0.5.

Finally, the harvest rate can be determined based on the knowledge of the feed and bleed rates such as to maintain a constant volume:

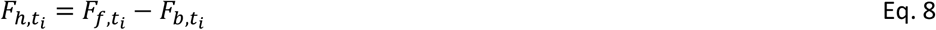

The perfusion culture was performed in 2L Univessel^®^ bioreactor with a working volume of 200 mL. The cultures were inoculated at 0.3×10^6^ cells/mL. The perfusion medium was a mix of 91.2% of PM, 8% FMA, 0.8% FMB and 6mM of L-glutamine. The cultures conditions were controlled at 36.8 °C for the temperature, 260 rpm for the agitation (adjusted to 300 rpm and 320 rpm during culture according O_2_ demand), pH 6.95 ± 0.05 with CO_2_ and 1M NaCO_3_, 60% of DO with O_2_ and air inlets. 1 ml of antifoam (Sigma antifoam C 2%) was automatically added every day and manually added if needed.

## 3 Theory/ Calculation / Modeling / Theoretical aspects

### 3.1 Model development

The model consists of a set of ordinary differential equations (ODEs) describing the dynamic of the population of cells as they move through three phases: live cells, dead cells, and lysed cells (Eq 9-11).

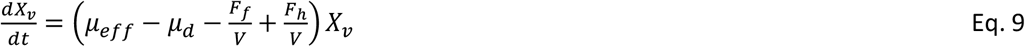

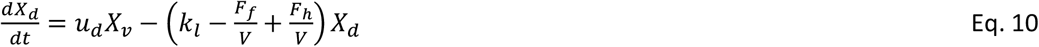

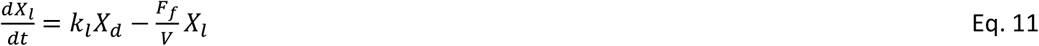

where *X_v_* is the viable cell density (VCD - concentration of viable cells), *X_d_* is the dead cell density (concentration of dead cells), and *X_l_* is the lysed cell density (concentration of lysed cells). *F_b_* is the bleed rate, *F_h_* is the harvest rate, and *V* is the reactor volume. *μ_eff_*, *μ_d_*, and *k_l_* are the effective growth, effective death, and lysing rates respectively.

The model also includes a catch-all “biomaterial” variable (∅*_b_*) representing a bulk set of metabolic byproducts secreted by the cells:

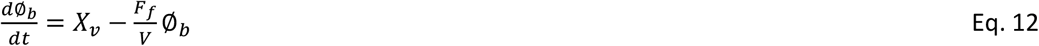

These mass balance equations are developed under the assumption that the bleed stream (*F_b_*) has the same content as the bioreactor and the harvest stream (*F_h_*) is cell free, assuming an ideal separation filter where the lysed cells and biomaterial pass through, and only viable and dead cells are retained into the bioreactor (Figure 1).

The cell growth rate (*μ_eff_*) is represented as the product of the maximal growth rate (*μ_max_*) and a nonlinear factor that describes the inhibition of growth due to the accumulation of byproducts (represented by the biomaterial variable ∅*_b_*). The resulting effective growth rate is captured in the following equation:

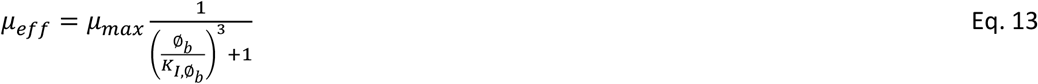

where *K*_*I*,∅_*b*__ is a parameter that represents the concentration of biomaterial ∅*_b_* above which inhibition occurs (Supp Figure 1).

The effective death rate, *μ_d_*, is dependent on a base death rate and a toxicity factor related to the accumulation of lysed cells (*X_l_*). Functionally:

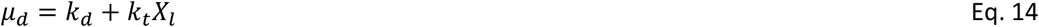

where *k_d_* is the primary death rate and *k_t_* represents the toxicity rate associated to the accumulation of lysed cells in the bioreactor.

Finally, the lysing process is governed by *k_l_* through a first-order rate law. Lysed cell material can exit the reactor by either the harvest stream or the bleed stream. Tracking the material balance of viable and dead cells gives an indication of total cells generated, and by extension the number of cells that have lysed and are no longer detectable.

Dead cells amount is evaluated indirectly through cell viability measurement which captures the ratio between viable cells and total cells:

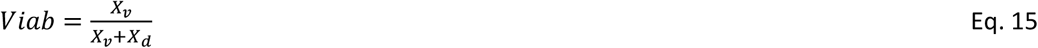

### 3.2 Parameter identification

Dynamic equations were solved by MATLAB’s ordinary differential equation solver function ode15s. The parameter identification was performed by using the Nelder–Mead simplex optimization algorithm (function *fminsearch*) in order to minimize a least-squares criterion (sum of squared differences between model predictions and experimental measurements).

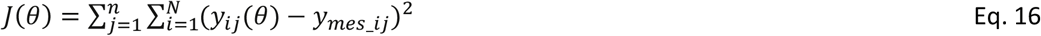

where *θ* is the vector of the parameters to be identified (dim *θ* = 5), 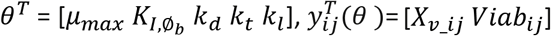 is the vector of the simulated variables (using model of mass balance equations 9–12) at the *i*^th^ time instant in the *j*^th^ experiment, 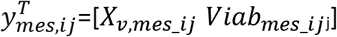 is the vector of the corresponding measurements.

### 3.3 Parameter sensitivity analysis and predicted model output uncertainty

The analysis of the sensitivity of the model outputs with respect to the parameters was performed as in Richelle et al.^15^. To this end, the four state variables (*X_v_*, *X_d_*, and ∅*_b_*) were defined as the system outputs *y_j_* with i = 1 : 4. The parameters were denoted *θ_j_* with j = 1 : 5. The time evolution of the 4 × 5 sensitivity functions 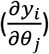 was then computed as follows:

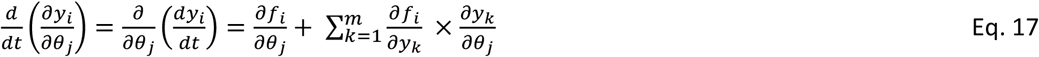

for i = 1 to 4, j = 1 to 4 and m = dim(y) = 4 with *dy_i_/dt = f_i_ (y, q_j_, t*) represented by model equations 9 - 12.

These sensitivity functions were used for computing a lower bound of the variance (Cramer-Rao bound) of the parameter estimation errors 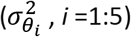 on the basis of the Fischer information matrix:

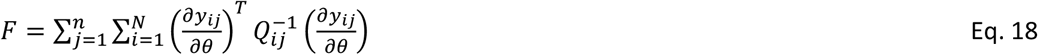

where 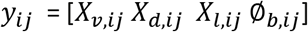 at the i^th^ time instant in the j^th^ experiment and 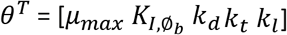.

The covariance matrix S could also be used to measure the correlation between the parameters (linear dependencies):

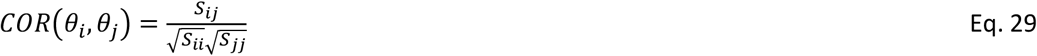

where *S_ij_* is the covariance of the errors on parameter estimates *θ_i_* and *θ_j_*; *S_ii_* and *S_jj_* are respectively the variance of the errors on parameter estimates *θ_i_* and *θ_j_*.

For analysing the uncertainty on the model outputs with respect to the parameter estimation errors, a global approach based on Monte Carlo sampling method was used. Contrarily to local approach based on first-order Taylor series approximation, this approach does not assume that the model responds linearly to a perturbation evaluated at a specific point of the parameter space. Instead, this sampling-based method uses a repeated random sampling of parameter values in a defined parameter space. In doing so, the overall model is used to generate the associated predicted model outputs by an iterative process of model simulations.

## 4 Results

### 4.1 Model identification using fed-batch cultures

We developed a growth model that tracks density and viability of a cell culture population (live, dead and lysed). The parameters of this model were identified based on 4 replicate fed-batch experiments performed in Ambr^®^ 250 (see Methods for details). To circumvent local minima and convergence problems with the optimization algorithm, a multi-start strategy was considered for the initialization of the parameter values. 100 uniformly distributed pseudo-random values over a given range (Table 1) were used for the initialization of the algorithm. For analysing the uncertainty on the model outputs with respect to the parameter estimation errors, a global approach with a Monte Carlo simulation was used, based on 1000 normally distributed pseudo-random sets of parameter values (Figure 2). In order to ensure the covering of the parameter space, the range (variability) for each parameter was determined by the confidence intervals presented in Table 1.

**Table 1.** Parameters values identified for each experiment separately and whole set of experiments 1 to 4

| | Initialization range | Exp. 1 | Exp. 2 | Exp. 3 | Exp. 4 | Exp. 1-2-3-4 | $\sigma_\theta^*$ | CV** |
| --- | --- | --- | --- | --- | --- | --- | --- | --- |
| $\mu_{max}$ | [0.01, 1] | 0.8409 | 0.8560 | 0.8145 | 0.8439 | 0.8384 | 0.0005 | 0.06 |
| $k_d$ | [0.01, 1] | 0.0210 | 0.0275 | 0.0273 | 0.0136 | 0.0209 | 0.0002 | 0.87 |
| $k_t$ | [0.01, 1] | 0.0286 | 0.0261 | 0.0232 | 0.0376 | 0.0290 | 0.0002 | 0.62 |
| $K_{I,\phi_b}$ | [1, 100] | 24.1117 | 24.4954 | 25.9156 | 23.7059 | 24.3905 | 0.0226 | 0.09 |
| $k_l$ | [0.01, 1] | 0.8723 | 0.7209 | 0.8765 | 0.6352 | 0.7743 | 0.0702 | 9.06 |

\*Standard deviation of parameter values identified using the whole set of experiments
\*\* Coefficient of variation (CV) of parameter values ( $\sigma_\theta/\theta$ - expressed in %) identified on the whole set of experiments 1 to 4

**Figure 2.**
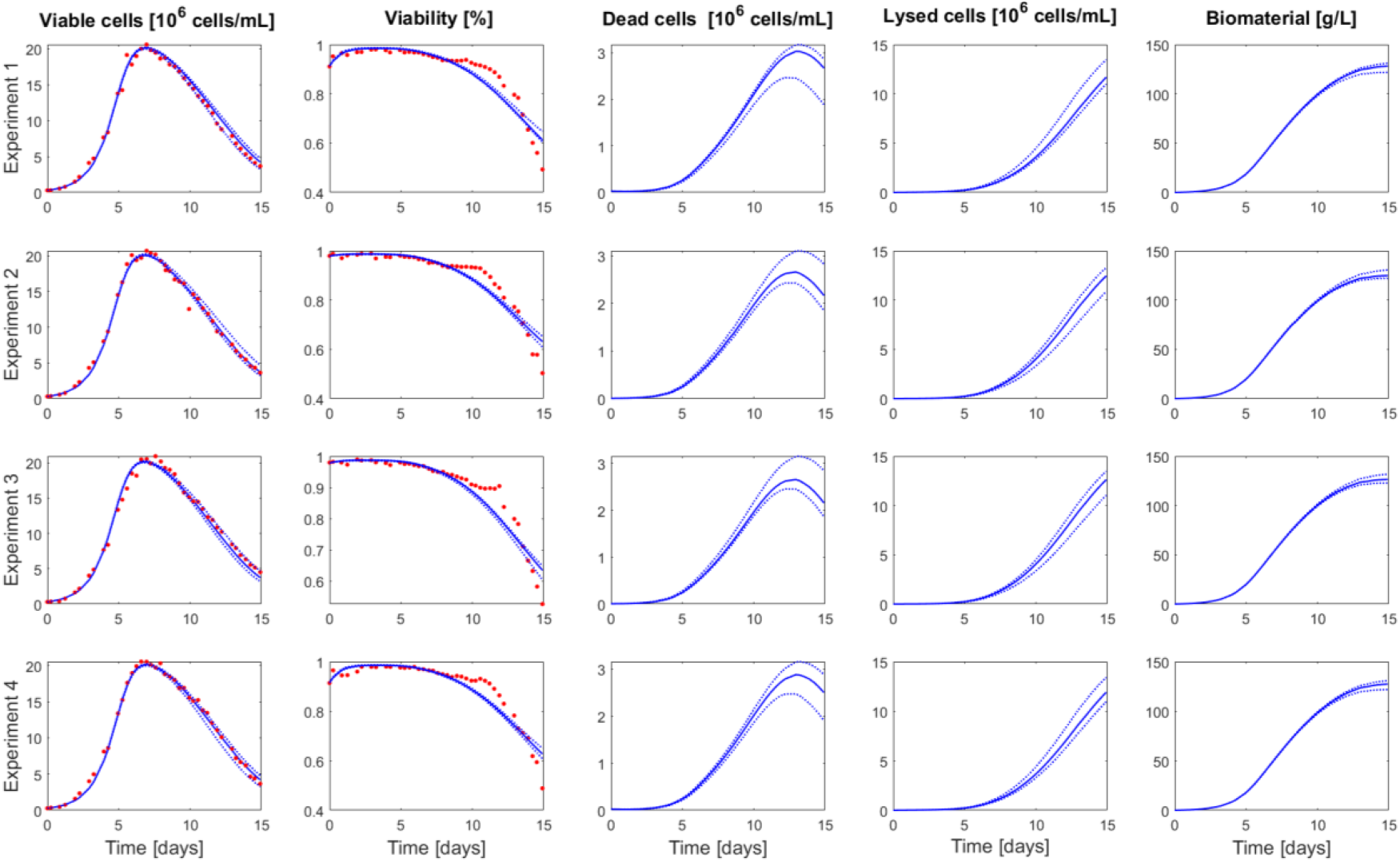
Comparison between measurements of Ambr^®^ 250 fed-batch experiments 1-4 (red dots) and the model simulation (blue curve) performed using the parameters value identified on the whole set of experiments. The dashed blue lines represent the uncertainty in the model predictions – calculated using Monte Carlo simulations (1000 samples) of normally distributed pseudo random parameters values (parameter space defined by *θ* ± 2*σ_θ_*)

The identified parameter values (based on the 4 experiments) are presented in Table 1 and the correlation matrix (absolute values of the correlation coefficients between parameters) in Table 2. Results obtained for the parameter identification of each experiment separately are also presented in Supplementary Tables 1-6. The model simulations and associated confidence intervals are presented in Figure 2 and Supplementary Figure 2 along with the experimental data used to identify the model.

**Table 2.** Correlation matrix (absolute value) of the parameters identified on the whole set of experiment

| | $\mu_{max}$ | $k_d$ | $k_t$ | $K_{I,\phi_b}$ | $k_l$ |
| --- | --- | --- | --- | --- | --- |
| $\mu_{max}$ | 1 | 0.1669 | 0.0626 | 0.6263 | 0.0415 |
| $k_d$ | 0.1669 | 1 | 0.7480 | 0.0731 | 0.1358 |
| $k_t$ | 0.0626 | 0.7480 | 1 | 0.0884 | 0.4355 |
| $K_{I,\phi_b}$ | 0.6263 | 0.0731 | 0.0884 | 1 | 0.0619 |
| $k_l$ | 0.0415 | 0.1358 | 0.4355 | 0.0619 | 1 |

The model captures well the dynamic of cell growth and the decrease over time of cell viability measurements for the 4 experiments. The parameters were identified with good confidence, this was also reflected in the simulation of the model output uncertainty (Table 1 and Figure 2). The largest uncertainty was associated to the lysed cells with the parameter *k_l_*. This is explained by the fact that lysed cells state variable is a degree of freedom for the model as no measurement are available. The highest parameter correlation is observed between *μ_max_* and 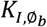 followed by *k_t_* and *k_l_* as expected due to their respective formulation of the effective growth and death rates (Eqs 13 and 14).

### 4.2 Model-based prediction of intensified operations performance

The model was further cross-validated using data from intensified cultures also performed in Ambr^®^ 250 (Experiments 5, 6 and 7 – see Methods for details). The model simulation successfully predicted the culture dynamic when transferred in intensified operations with media exchange. For these intensified cultures, the culture media was harvested at the same rate as the medium feeding; keeping the lived and dead cells into the bioreactor while lysed cells and secreted biomaterials were removed thanks to the presence of a cell retention device (Figure 1). Doing so, the growth was no longer inhibited by the accumulation of by-products (represented with the biomaterial variable) and the death rate was less favored by the accumulation of the lysed cells in the media. Specifically, the biomaterial concentration of lysed cells and biomaterials after 10 days of culture in intensified conditions were respectively 10- and 4-fold lower than for fed-batch operations (Figure 3).

**Figure 3.**
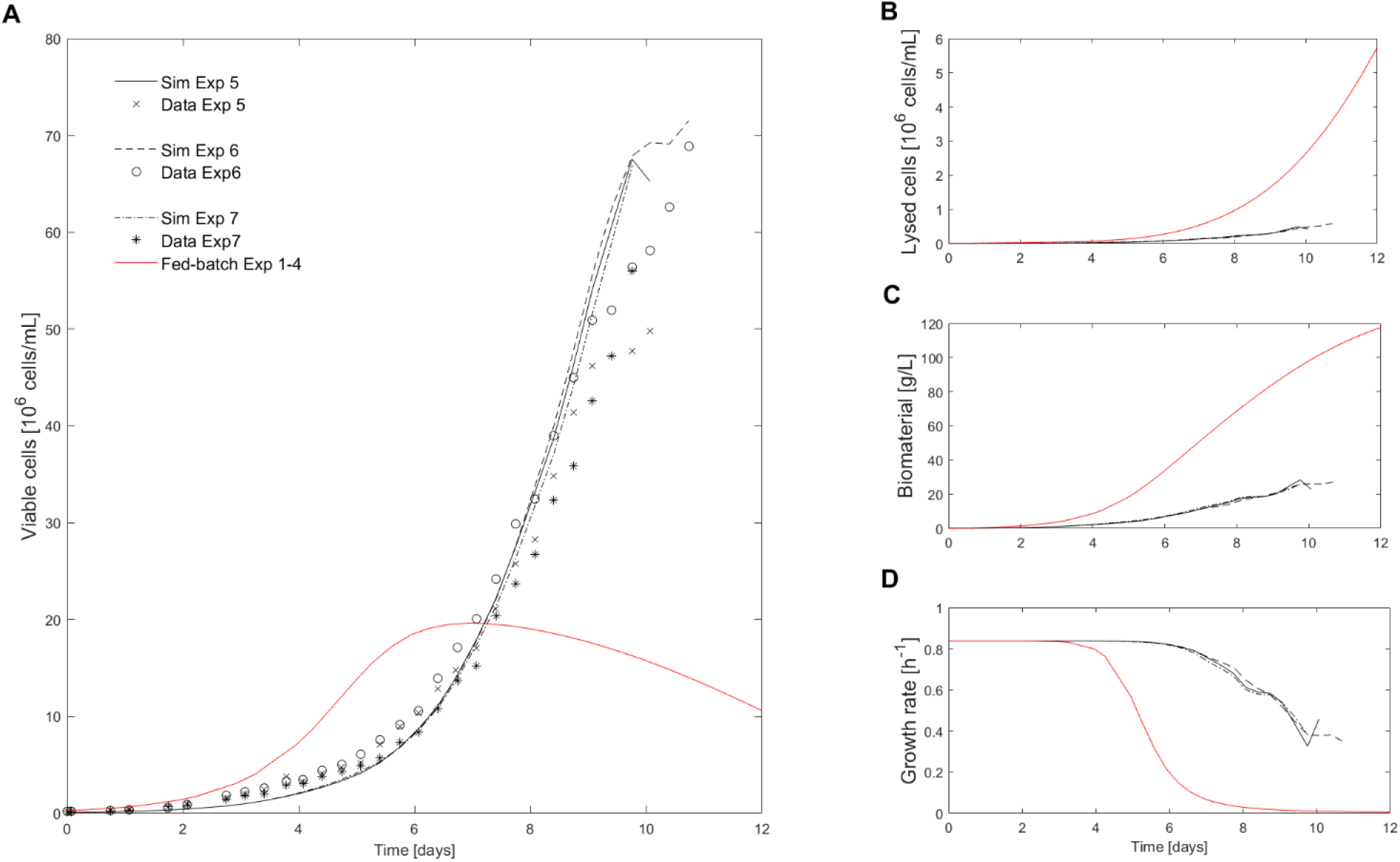
Comparison of intensified (media exchange) and fed-batch experiments. A. Comparison of viable cell density measurement of intensified cultures 5-7 (black cross, star and open circle) with the model simulation of the intensified culture (solid black line) and the fed-batch culture (solid red-line) performed using the parameters value identified on the whole set of fed-batch experiments 1-4. B, C and D present, respectively, a comparison of the lysed cells density, biomaterial concentration and growth rate simulated for the intensified culture (solid black line) and the fed-batch culture (solid red-line) using the parameters value identified on the whole set of fed-batch experiments 1-4

The proposed model has a rather simple structure compared to the ones presented in literature. The overall formalism to describe the different states of cells is conserved across existing models: cell growth and mortality occurs in parallel while dead cells are lysed over time. The main difference in our proposed structure lies in the description of the growth and dead rates. Indeed, it is well known that mammalian cell metabolism can be limited by the depletion of nutrients or by the accumulation of inhibitory metabolites^16^. Therefore, the death and growth rate are typically described as extended Monod’s law (more than one compound influence the reaction rate) accounting for diverse activating and inhibiting compounds.

For example, Shirahata et al.^17^ modelled the growth rate in continuous operation with an inhibition by the accumulation of ammonia. Lourenço da Silva et al.^16^ developed a kinetic model that describes the growth of hybridoma cells in fed-batch culture with decreasing and death enhancing effects of glucose, amino-acids, serum and oxygen depletion, on the one hand, and of ammonia and lactate accumulation on the other. Craven et al.^18^ accounted in their growth model for the activation by substrates (glucose and glutamine) and inhibition by by-products (lactate and ammonia). Papathanasiou et al.^13^ used five metabolites (glucose, glutamine, arginine, aspartate, asparagine) to describe their activation and inhibition influence the respective growth and death process.

The evaluation of the respective influence of these potential limiting factors is a difficult task as several of these factors are often simultaneously limiting, leading to observed diversity in the model formalism for growth and death rates. Furthermore, the description of such activation and inhibition effects by multiple metabolites quickly complicates the model structure. Indeed, with this formalism, these compounds are introduced as state variables in the model and their associated kinetics need to be described. The main novelty of the proposed model is the introduction of a catch-all “biomaterial” variable. This variable captures the inhibition of growth by a bulk of secreted by-products without detailing the identity and contribution of each potential inhibitor. Therefore, it simplifies the model structure (and reduced the number of model parameters) as there is no need to describe the dynamic associated to these compounds.

### 4.3 Design and analysis of perfusion process conditions

The expected growth profile in a perfusion operation mode was simulated using the following assumptions:

- The simulation was performed for a 2L bioreactor Univessel^®^
- Initial seeding density and viability was set to the same from as the Ambr^®^ 250 fed-batch experiments
- The lysed cells and inhibitory biomaterial were initialized at 0
- There were no considerations for adjustment in growth changes during the simulation
- It was assumed that the media composition and perfusion rate is sufficient for supplying nutrients

Different events for process operation changes were introduced to test the capabilities of the model and the cell’s response to switch in operations (Table 3):

- The culture began with an intensification phase to reach the cell density target (*X_V,target_*= 50. 10^6^ cells/mL). The feed and harvest rate were equal (*F_f_*=*F_h_*) and defined as presented in Methods for a perfusion rate (*P*) of 2.25 vol/day. The bleed rate (*F_h_*) was equal to zero
- Once the cell density reached 95% of the desired target *X_V,target_*, a PI controller was used for adjusting the bleed rate and maintaining a desired setpoint (details of the PI control setup presented in Methods)
- An increase in the perfusion rate (*P*) was introduced for more than a day to test the PI control (in between 12,9 and 14.1 days) before being set back to its original defined value of 2.25 vol/day
- A decrease of the perfusion rate (*P*) was imposed at 21.1 days to assess the response of the cells to an increase of biomaterial accumulation
- Finally, an increase of the cell density target (to *X_V,target_*= 70. 10^6^ cells/mL) was introduced to evaluate the capacity of the cells to cope with strong switch in operations

**Table 3.** Details of switch in process operations for perfusion simulation and experimental run.

| Event time [days] | $P$ [vol/day] | $X_{V,target}$ [$10^6$ cells/mL] |
| --- | --- | --- |
| 0 | 2.25 | 50 |
| 12.9 | 3 | 50 |
| 14.1 | 2.25 | 50 |
| 21.1 | 1.75 | 50 |
| 24 | 1.75 | 70 |

The simulation of this perfusion experiment was presented in Figure 4 and Supplementary Figure 4 along with the experimental data collected during a 2L perfusion bioreactor run performed under the same operations listed in Table 3. The model prediction accurately captured the dynamic of the cell culture in perfusion based on parameter values identified using Ambr^®^ 250 fed-batch experiments. The model also accurately identified the decrease of cell viability initiated once the perfusion rate was decreased (*P* =1.75 vol/day) due to the accumulation of biomaterials and the maximum stable target cell density at the last process operations switch (*X_V,target_* = 70. 10^6^ cells/mL). The PI controller adequately adjusted the bleed rate to hold a stable cell density (simulations were in agreement with stream rates implemented by the online PI controller – Supplementary Figure 3).

**Figure 4.**
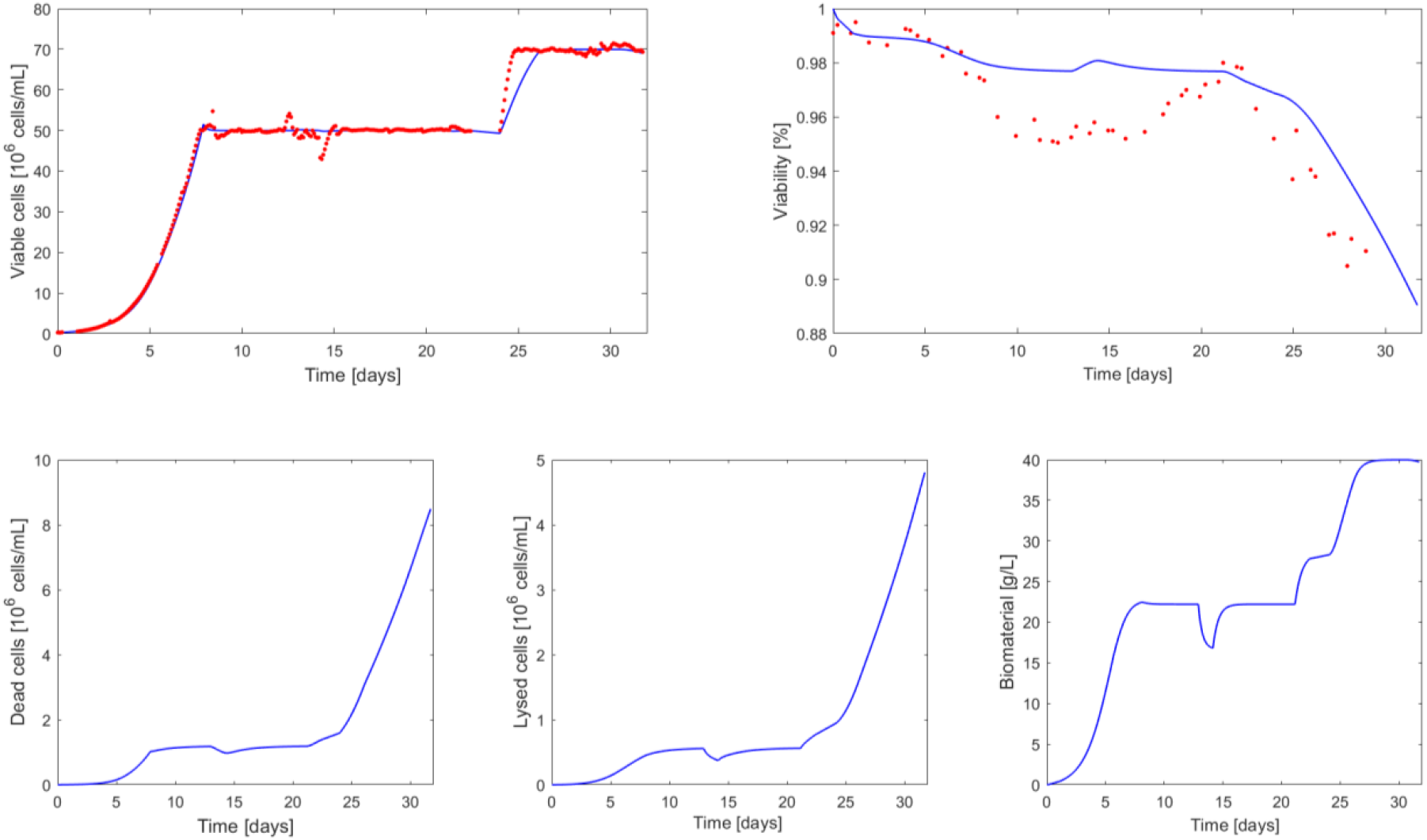
Comparison between measurements of 2L perfusion experiment 8 (red dots) and the model simulation (blue curve) performed using the parameters value identified on the whole set of fed-batch experiments 1-4.

To our knowledge this is the first design of a perfusion culture using a model identified based on fed-batch experiments. Typically, models are developed for one type of culture (batch, fed-batch or continuous) and cannot be transferred to other process operation without changes in the model structure and/or parameter values. Shirahata et al.^17^ modified the formalism of the growth rate function depending on the operation mode. Specifically, in batch mode, they simulated the viable cell dynamic using an activation by glucose and the onset of massive cell death when a glucose depletion occurs. In perfusion mode, the growth rate was no longer modelled in function of substrate consumption but rather with an inhibition due to ammonia accumulation in the culture medium. Lourenço da Silva et al.^16^ successfully validated a kinetic model for hybridoma fed-batch culture and mentioned that they were capable to simulate experimental results obtained during batch and continuous processes with minor changes of few kinetic parameters. Unfortunately, the data were not shown. Finally, Craven et al.^18^ developed a unique model structure for CHO cell culture operated under 3 different modes (batch, bolus fed-batch and continuous fed-batch) and grown under 2 scales (3 and 15 L) but the model parameters identified changed with scale and mode of operation.

The presented model-based approach represents a reliable alternative to existing experimental procedure such as the ones presented in Janoschek et al.^19^ and Wolf et al.^20^. These protocols rely on the evaluation and optimization of different feed, harvest and bleed strategies similar to a Design of Experiments (DoE) approach. While these methods have been proven to be successful, they are experimentally intensive and do not allow the user to test the system response to joint variation of multiple control variables and setpoints.

## 5 Conclusions

The era of digital transformation has reached biopharma, and the companies that will find success in the next phase will be those who adopt innovative approach to accelerate product development and production. In this context, in-silico computational tools to help optimize upstream bioprocesses will be essential^21^.

The goal of this study was to propose a model-based strategy to improve upstream cell culture development within biopharmaceutical manufacturing thanks to its process transfer capabilities. Often referred to as in-silico experimentation, subject matter experts (SME) can use the proposed framework to digitally test various hypothetical operating policies. Ideas can be honed and proposed before verifying in the lab. The hypothesized model was built from limited data with a focus on core growth kinetics and sensitivity to biomaterials. It can be used to investigate growth trajectories and evaluate media exchange operating modes (intensified growth and perfusion). To demonstrate these capabilities, the model was calibrated with Ambr^®^ 250 fed-batch experiments and successfully used to forecast growth profiles under various operating modes including the cell line’s response to media exchange.

As more experiments are run and data is collected, this generic model structure can continuously be extended to include additional metabolic information from shifts in pH, temperature, media composition and other important process conditions. Such model would therefore also be used to optimize media composition and recipe decisions to maximize productivity while maintaining Critical Quality Attributes (CQAs) within specification.

However, models describing the influence of spent media composition on productivity and product quality are far more complex and, currently, not as mature as those for growth description. This relates to growth and death kinetics being driven strongly by the extracellular environment, while productivity and CQAs (e.g., glycan profile) are influenced through more subtle shifts in the intracellular metabolism. For the moment, intracellular measurements are expensive and not practical for typical product development workflows or high throughput experimental designs. Therefore, analytical tool such as machine learning and other data driven methods would most likely be used to relate extracellular process measurements to the CQAs.

To conclude, using simulation is common practice in many process industries but a relatively new tool for biopharmaceutical manufacturing. Being able to test operating strategies digitally reduces wet lab experimental needs, speeding up the product development process. The big picture then – and the takeaway for biopharma companies – is to move toward an enhanced, optimized approach to upstream process development that makes use of existing information to bring transformation, optimization and ultimately, profitability. The key dynamic behind all of it is an integration of advanced data analytics, process knowledge and digital tools that transcend the traditional method of process monitoring and move toward digital twins powered by a systems approach of bio-simulation.

## Acknowledgements

We thank Steffi Scholze and others in the Sartorius Corporate Research team for their support in in the development of this study. We also thank Timo Schmidberger and the Sartorius Data Analytics team for their investment in the model evaluation, validation and benchmarking. The US Food and Drug Administration for their support in the development of in-silico process design and optimization strategies. Our collaborators at GSK and Sanofi for testing the proposed model-based process design strategy.

## Notes

### Competing Interest Statement

The authors have declared no competing interest.

