## Supplementary material for "Model-based intensification of CHO cell cultures: one-step strategy from fed-batch to perfusion"

Supp. Table 1 - Standard deviation of parameter value identified for each experiment separately

| | Exp. 1 | Exp. 2 | Exp. 3 | Exp. 4 |
|---|---|---|---|---|
| $\mu_{max}$ | 0.0010 | 0.0010 | 0.0010 | 0.0010 |
| $k_d$ | 0.0005 | 0.0004 | 0.0003 | 0.0003 |
| $k_t$ | 0.0004 | 0.0003 | 0.0007 | 0.0007 |
| $K_{I,\emptyset_b}$ | 0.0459 | 0.0437 | 0.0465 | 0.0444 |
| $k_l$ | 0.2132 | 0.3703 | 0.1043 | 0.0670 |

Supp. Table 2 - Coefficient of variation (CV) of parameter values ( $\sigma_\theta/\theta$ - expressed in %) identified for each experiment separately

| | Exp. 1 | Exp. 2 | Exp. 3 | Exp. 4 |
|---|---|---|---|---|
| $\mu_{max}$ | 0.12 | 0.12 | 0.13 | 0.12 |
| $k_d$ | 2.18 | 1.29 | 2.49 | 2.14 |
| $k_t$ | 1.34 | 1.09 | 2.02 | 1.88 |
| $K_{I,\emptyset_b}$ | 0.19 | 0.18 | 0.19 | 0.19 |
| $k_l$ | 24.44 | 51.36 | 13.43 | 10.55 |

Supp. Table 3 - Correlation matrix (absolute value) of the parameters identified for Experiment 1

| | $\mu_{max}$ | $k_d$ | $k_t$ | $K_{I,\emptyset_b}$ | $k_l$ |
|---|---|---|---|---|---|
| $\mu_{max}$ | 1 | 0.1472 | 0.0678 | 0.6216 | 0.0428 |
| $k_d$ | 0.1472 | 1 | 0.8146 | 0.0620 | 0.2018 |
| $k_t$ | 0.0678 | 0.8146 | 1 | 0.0874 | 0.2770 |
| $K_{I,\emptyset_b}$ | 0.6216 | 0.0620 | 0.0874 | 1 | 0.0626 |
| $k_l$ | 0.0428 | 0.2018 | 0.2770 | 0.0626 | 1 |

Supp. Table 4 - Correlation matrix (absolute value) of the parameters identified for Experiment 2

| | $\mu_{max}$ | $k_d$ | $k_t$ | $K_{I,\emptyset_b}$ | $k_l$ |
|---|---|---|---|---|---|
| $\mu_{max}$ | 1 | 0.2089 | 0.0910 | 0.6209 | 0.0424 |
| $k_d$ | 0.2089 | 1 | 0.7794 | 0.0960 | 0.1142 |
| $k_t$ | 0.0910 | 0.7794 | 1 | 0.1303 | 0.3462 |
| $K_{I,\emptyset_b}$ | 0.6209 | 0.0960 | 0.1303 | 1 | 0.0615 |
| $k_l$ | 0.0424 | 0.1142 | 0.3462 | 0.0615 | 1 |

Supp. Table 5 – Correlation matrix (absolute value) of the parameters identified for Experiment 3

| | $\mu_{max}$ | $k_d$ | $k_t$ | $K_{I,\emptyset_b}$ | $k_l$ |
|---|---|---|---|---|---|
| $\mu_{max}$ | 1 | 0.1165 | 0.0029 | 0.6560 | 0.0351 |
| $k_d$ | 0.1165 | 1 | 0.4454 | 0.0489 | 0.1229 |
| $k_t$ | 0.0029 | 0.4454 | 1 | 0.0081 | 0.8097 |
| $K_{I,\emptyset_b}$ | 0.6560 | 0.0489 | 0.0081 | 1 | 0.0548 |
| $k_l$ | 0.0351 | 0.1229 | 0.8097 | 0.0548 | 1 |

Supp. Table 6 – Correlation matrix (absolute value) of the parameters identified for Experiment 4

| | $\mu_{max}$ | $k_d$ | $k_t$ | $K_{I,\emptyset_b}$ | $k_l$ |
|---|---|---|---|---|---|
| $\mu_{max}$ | 1 | 0.1538 | 0.0076 | 0.6444 | 0.0459 |
| $k_d$ | 0.1538 | 1 | 0.4376 | 0.0593 | 0.1163 |
| $k_t$ | 0.0076 | 0.4376 | 1 | 0.0046 | 0.8201 |
| $K_{I,\emptyset_b}$ | 0.6444 | 0.0593 | 0.0046 | 1 | 0.0675 |
| $k_l$ | 0.0459 | 0.1163 | 0.8201 | 0.0675 | 1 |

Supp Figure 1 - Growth inhibition factor ($f(\emptyset_b)$) as a function of the biomaterial concentration ($\emptyset_b$) value for a range of parameter values $K_{I,\emptyset_b}$.

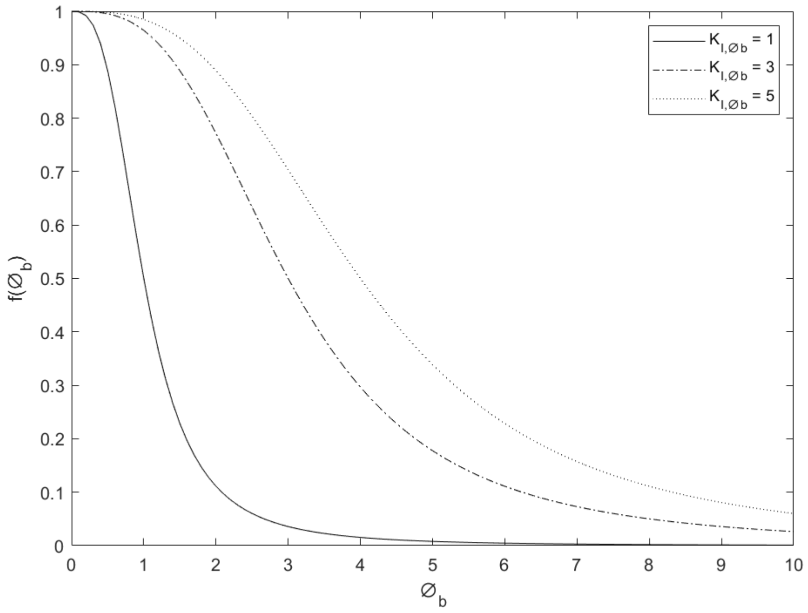


Supp. Figure 2 - Comparison between measurements of Ambr250 fed-batch experiments 1- 4 (red dots) and the model simulation (blue curve) performed using the parameters value identified for each experiment separately. The dashed blue lines represent the uncertainty in the model predictions – calculated using Monte Carlo simulations (1000 samples) of normally distributed pseudo random parameters values (parameter space defined by $\theta \pm 2\sigma_\theta$)

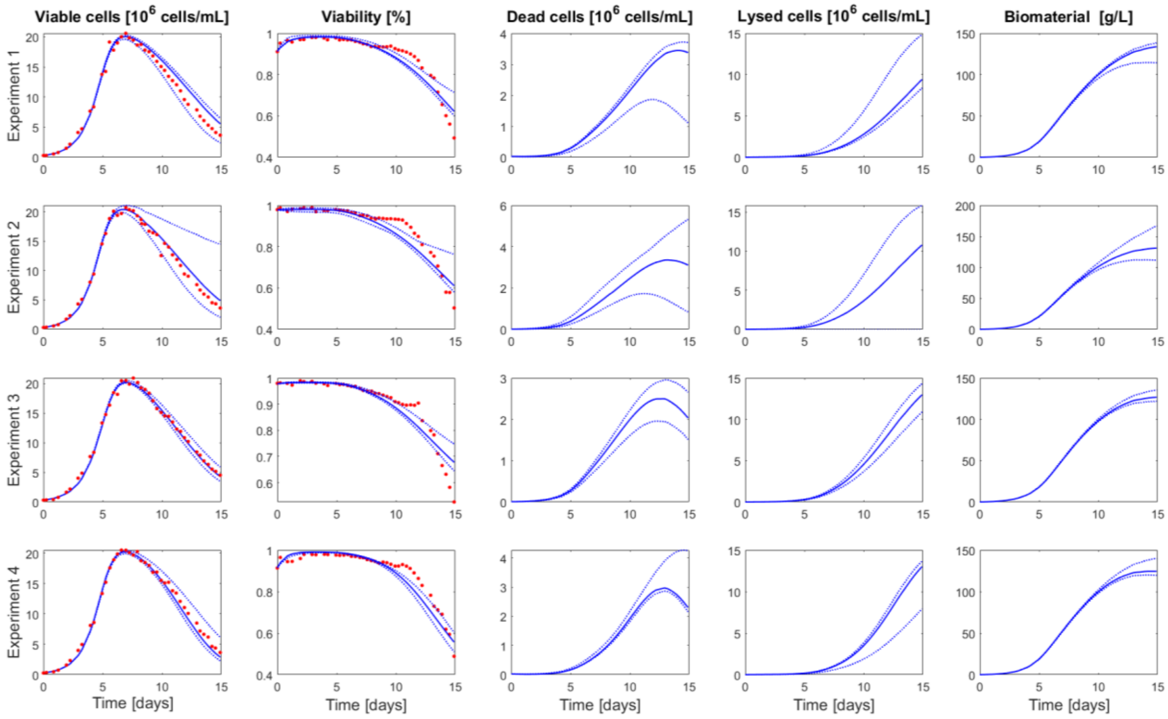

Supp. Figure 3 - Comparison between the feed, harvest and bleed rates implemented by the PI controller during 2L perfusion experiment 8 (red dots) and the model simulation (blue curve) performed using the parameters value identified on the whole set of fed-batch experiments 1-4.

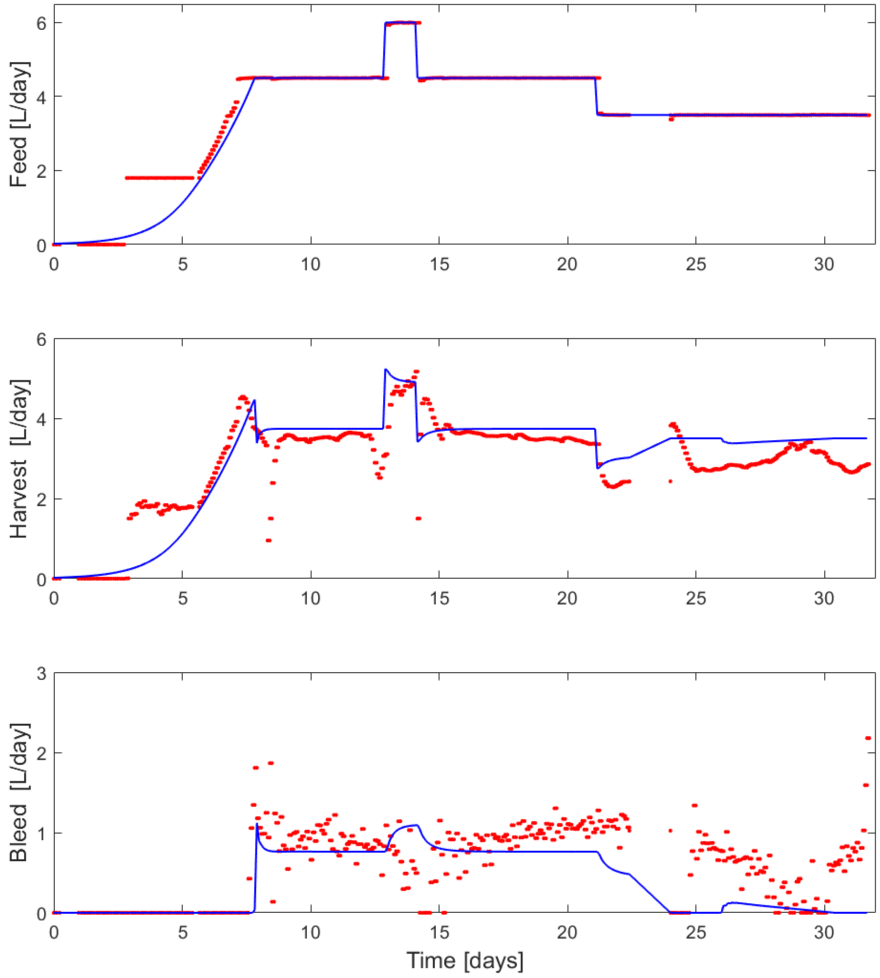
